# Ouro-seq: Improved Recovery of Full-Length circRNAs from Samples with Limited RNA Content

**DOI:** 10.64898/2026.08.14.744779

**Authors:** Birgit M.M. Wever, Yara van den Burgt, Florent Mouliere, D. Michiel Pegtel, Maaike C.G. Bleeker, Renske D.M. Steenbergen, Norbert Moldovan

**Author notes:** These authors contributed equally (Birgit M.M. Wever, Yara van den Burgt). Senior authors (Renske D.M. Steenbergen, Norbert Moldovan).

## Abstract

Circular RNAs (circRNAs) are an emerging class of RNAs with biomarker potential, but their detection in liquid biopsies is challenging due to low abundance. We developed Ouro-seq, a novel long-read sequencing protocol optimized for full-length circRNA recovery. Applied to urine, cervico-vaginal self-samples from cervical cancer patients, and plasma from lung cancer patients and controls, Ouro-seq recovered 2-5 times more and substantially longer circRNA molecules than conventional methods. Plasma contained predominantly exonic circRNAs, while urine and cervico-vaginal samples were dominated by previously undercharacterized intergenic circRNAs. We also identified extensive alternative circularization and splicing events. Functional analysis revealed distinct specialization patterns: exonic circRNAs showed enhanced miRNA sponging potential, while circRNAs from unplaced genomic scaffolds demonstrated greater peptide-coding capacity. This study establishes Ouro-seq as a valuable tool for comprehensive circRNA characterization in low-yield clinical samples and advances circRNA biology understanding with potential biomarker discovery and disease monitoring applications.

**Motivation:** While circular RNAs (circRNAs) constitute a minor fraction of total RNA, they may play critical roles in cancer development. CircRNA concentrations are typically too low for detection by Oxford Nanopore Long-Read Sequencing (LRS), particularly in samples with limited RNA content, such as liquid biopsies. Consequently, LRS-based circRNA analysis from liquid biopsies remains unexplored. To overcome these technical limitations, we developed an optimized circRNA enrichment method utilizing short-amplicon suppression, enabling circRNA profiling from urine, plasma, and cervico-vaginal samples.

## Introduction

Circular RNAs (circRNAs) are an emerging class of cancer biomarkers due to their role in cancer development and metastasis.^1^ Unlike linear RNAs, circRNAs form covalently closed loops, resulting in increased resistance to degradation.^2–4^ This relative stability makes circRNAs particularly attractive as biomarker candidates for liquid biopsy-based non-invasive cancer detection and monitoring.

Traditional short-read sequencing (SRS) methods face challenges in capturing the structural complexity and expression profiles of RNA isoforms, including circRNA.^5,6^ These methods rely on detecting a singular unique feature of circRNAs known as the back splice junction (BSJ). Furthermore, due to sequencing length limitations, SRS-based approaches use a probabilistic model of overlapping reads to reconstruct potential circRNA isoforms.^6,7^

In recent years, long-read sequencing (LRS) technologies, such as Oxford Nanopore sequencing, have enabled the comprehensive profiling of full-length transcripts.^8–10^ LRS enables coverage of the entire length of circRNA molecules providing a distinct advantage for the detection of complex alternative circularization events and splice variants.^6,11,12^

Despite the potential of circRNAs as biomarkers, their scarcity relative to linear RNAs poses a significant challenge for LRS. Current circRNA enrichment strategies rely on enzymatic degradation of linear RNA and/or poly(A) purification prior to reverse transcription (RT). Several LRS methodologies exist to further enrich cDNA originating from circRNA. CIRI-long uses a long fragment selection before sequencing to further eliminate the short leftovers of linear RNA digestion.^6^ isoCirc creates a circular RT product that is further amplified using rolling-circle amplification.^11^ circNick-LRS fragments the circRNA-enriched RNA pool and then performs the RT and downstream amplification.^12^ circPanel-LRS is a targeted approach using a panel of circRNA-specific primers to enrich circRNA reads from total RNA.^12^ These methods remain limited due to low enrichment of total circRNA yields from tissue and cell culture, with 0.1–6% of sequenced molecules representing circRNA species, provide fragmented information about the molecule or only inform about specific circRNA.^6,11–13^ In liquid biopsies, these limitations are further aggravated by their limited RNA yield compared to tissue.^14^

We present Ouro-seq *(Ouro derived from the name of Ouroboros, a mythical snake eating its own tail, a reference to the panhandle formation of short cDNA molecules as key feature of our method)*, a novel workflow for sequencing full-length circRNAs from samples with limited RNA content, such as plasma, urine and cervico-vaginal self-samples (CVS) using Oxford Nanopore LRS technology. We demonstrate that Ouro-seq improves full-length circRNA recovery, resulting in enhanced structural profiling of circRNA isoforms uncovering novel circRNAs with potential clinical relevance compared to conventional Linear RNA Degradation Only (LRDO) and LRS sequencing.

## Results

### The Ouro-seq protocol

To enrich for circRNA, we developed a customized approach using Nanopore amplified cDNA-seq library preparation (**Fig. 1A**). In brief, following total RNA extraction we performed ribosomal RNA (rRNA) depletion and linear RNA degradation. Reverse transcription (RT) was performed with a custom cDNA RT adapter composed of a short-amplicon suppression adapter and a random-hexamer 3’ overhang, followed by second-strand synthesis using a custom strand-switching primer containing the suppression adapter. This resulted in long concatemeric cDNA originating from circRNA and short cDNA from the fragments left over after linear RNA digestion. Short amplicon suppression PCR was performed on the cDNA with a single primer, during which inverted terminal repeats of the suppression adapters (**Supplementary information 1**) hybridize on shorter fragments blocking the single primers (**Fig. 1A step 4**). This resulted in a decreased PCR efficiency of shorter cDNA originating from linear fragments. After a final magnetic bead-based selection for long amplicons, barcoded Nanopore sequencing libraries were prepared and sequenced using the PromethION platform (see Methods).

**Figure 1.**
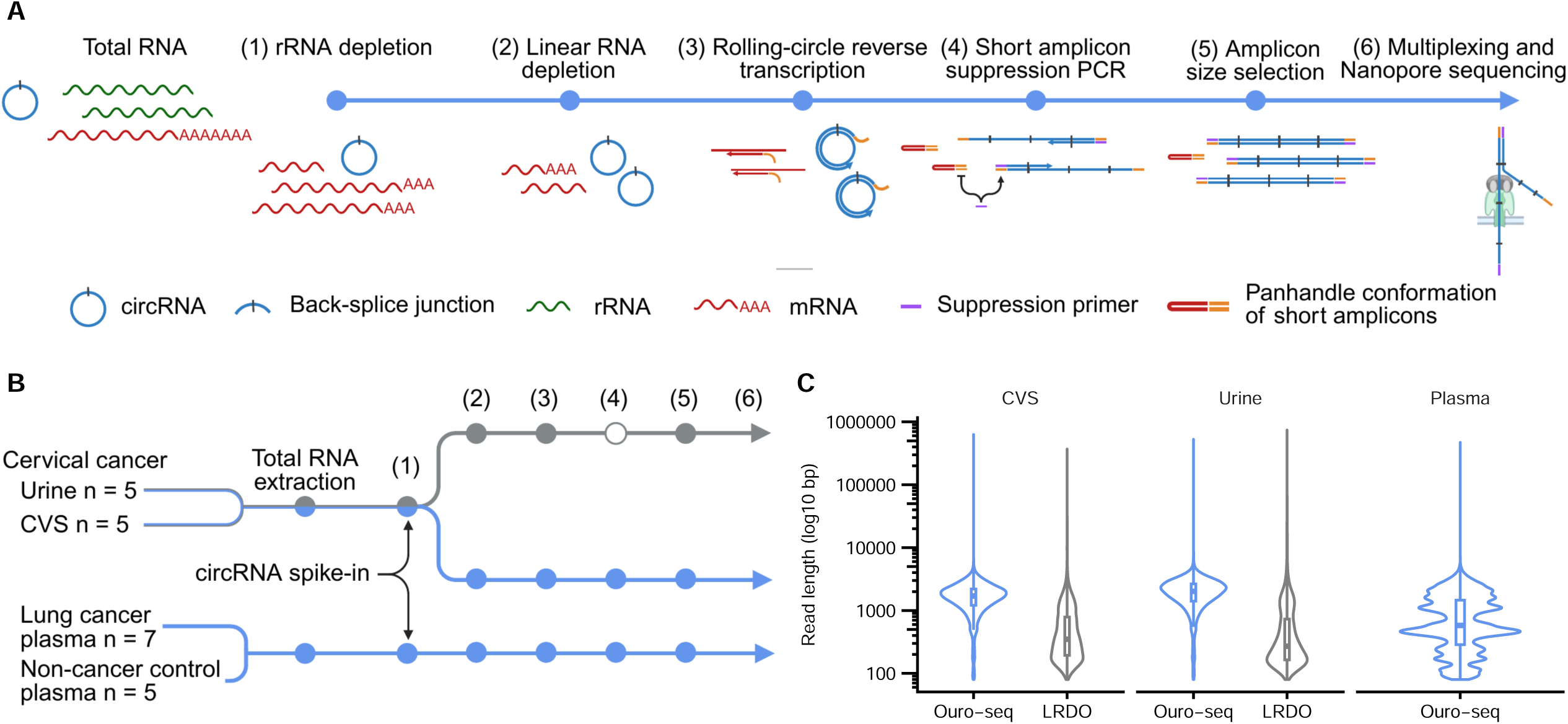
The Ouro-seq protocol. **A.** Schematic diagram of the Ouro-seq library preparation process. **B.** Schematic view of experimental conditions. The gray line: Linear RNA Degradation Only (LRDO) library preparation, blue line: Ouro-seq. (1) rRNA depletion, (2) linear RNA depletion, (3) Rolling-circle reverse transcription, (4) Short amplicon suppression PCR, (5) Amplicon size selection, (6) Multiplexing and Nanopore sequencing. Filled circles: library preparation steps. Empty circle: library preparation step not in LRDO.

Total RNA quantification confirmed anticipated low RNA concentrations in liquid biopsy, including urine (44.52 ± 38.62 ng/μL, mean ± SD) and plasma (8.00 ± 0.83 ng/μL). In contrast, CVS exhibited substantially higher RNA concentrations (232.42 ± 215.13 ng/μL), likely attributable to cellular contribution in CVS (**Table S1**). Following rRNA depletion and linear RNA degradation we were left on average with 5% of the starting RNA amounts with a mean of 9.4 ± 5.79 ng/μL for CVS and a mean of 0.33 ± 0.13 ng/μL for plasma. Urine sample measurements were excluded from this analysis due to inconsistencies—one sample exhibited concentrations exceeding total RNA values while two samples could not be quantified due to the device’s inability to detect internal control markers, despite visible peaks in the electropherogram (**Table S1**).

The Ouro-seq protocol resulted in a mean of 3,899,270 ± 3,046,915 sequencing reads for urine samples, a mean of 4,492,071 ± 5,833,118 sequencing reads for CVS and a mean of 1,378,307 ± 476,399 sequencing reads for plasma samples (**Table S2**).

We directly compared Ouro-seq with conventional LRDO long-read sequencing (**Fig. 1B**, see Methods) using the same urine and CVS samples. Plasma circRNA yield was too low to include in this direct comparison. The LRDO protocol resulted in 97,456.4 ± 29,492.24 reads for the urine and 91,097 ± 25,424.83 reads for the CVS samples (**Table S2**). Ouro-seq on average yielded longer reads compared to LRDO sequencing for both the urine (2,043.53 ± 1,020.46 bp versus 869.53 ± 7,258.15 bp) and CVS (1,738.02 ± 874.99 bp versus 742.32 ± 3,792.69 bp), and for plasma Ouro-seq resulted in a read length of 956.18 ± 913.61 bp (**Fig. 1C**, **Table S2**). These results show that Ouro-seq increases the sequencing yield and is selective for longer reads.

### Ouro-seq improves full-length circRNA recovery

We used the CIRI-long tool for circRNA calling and isoform detection.^6^ We defined full-length circRNA as cyclic consensus sequences (CCS) with a BSJ flanked by a canonical splice signal (AG/GU) or a known alternative splice signal (AG/GC, AC/AU, AC/GU or AG/AU).^6^ For urine samples, Ouro-seq reached on average a 4.94 times higher (from 0.034% to 0.17% of total reads, p = 0.008), for CVS on average a 2.18 times higher (from 0.062% to 0.14% of total reads, p = 0.03) circRNA yield compared to the LRDO method, while for plasma the full-length circRNA proportion was 0.25% of the total reads (**Fig. 2A**). To show that our method is not limited to low circRNA levels, we spiked in a synthetic circular molecule after total RNA extraction at equal concentration (see Methods, **Fig. 1B**). Ouro-seq enriched the spike-in on average 501.9 times (from 0.0032% to 1.62%, p = 0.05) in urine and on average 168.71 times (from 0.018% to 3.1%, p = 0.008) in CVS samples compared to the LRDO protocol, while the contribution of spike-ins in plasma samples was on average 69.95% (**Fig. 2B**). Ouro-seq detected longer circRNA with a length of 1,308.17 ± 835.89 bp compared to the LRDO method with a length of 62.39 ± 17.98 bp (p < 0.0001) for urine and 863.7 ± 608.55 bp compared to 66.91 ± 64.78 bp (p < 0.0001) in CVS, while plasma had a circRNA length of 2,991.65 ± 4,985.34 bp (**Fig. 2C**). Ouro-seq performs significantly better than LRDO for full-length circRNA recovery both for urine and CVS, however its performance falls behind for plasma, presumably due to the overrepresentation of spike-ins (**Fig. 2D**). These results show that Ouro-seq improves full-length circRNA recovery and is suitable for exploring long circRNA. They also confirm that plasma, urine and CVS samples have a low proportion of circRNA.

**Figure 2.**
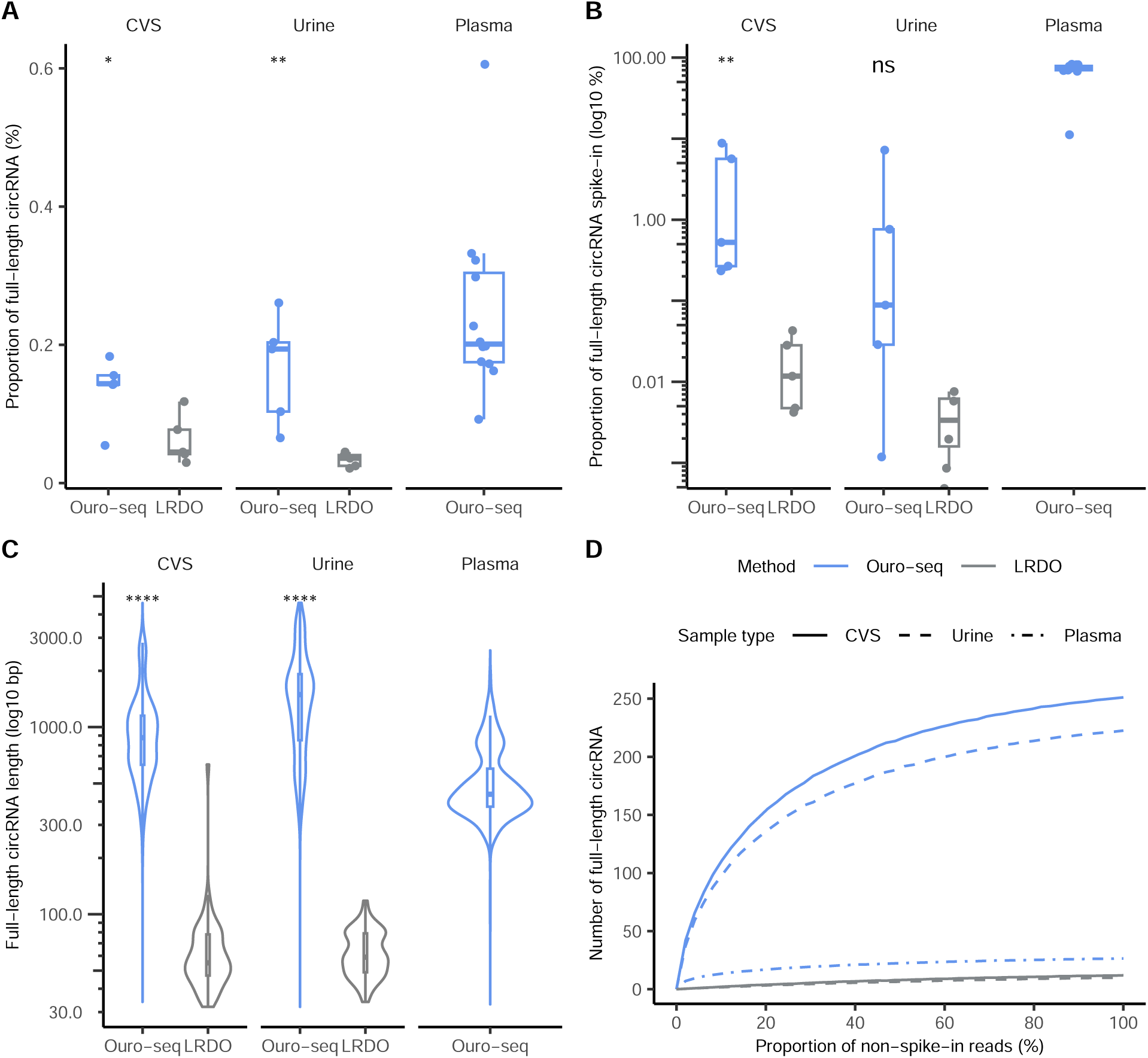
Ouro-seq improves full-length circRNA recovery. **A.** Proportion of full-length circRNA. **B.** Proportion of full-length circRNA spike-in. CVS: n = 5, urine: n = 5, plasma: n = 12. **C.** The length of full-length circRNA. circRNA counts weighted with read counts. Statistical significance assessed using the Mann-Whitney U-test: ****: p<0.0001, **: p<0.01, *: p<0.05, ns: not significant. **D.** Saturation curves of full-length circRNA detections. For each library preparation type and sample type reads were randomly sampled from real data in 100 replicates and the corresponding read count and full-length circRNA counts were calculated.

### Ouro-seq uncovers circRNA complexity

Ouro-seq detected a total of 1,111 circRNA in urine, 1,288 circRNA in CVS and 176 circRNA in plasma samples confirmed by at least two reads (**Table S3**). One hundred and two circRNA were detected in both urine and CVS. One hundred and fifty one circRNA from plasma, while only 2 from urine and 7 from CVS could be detected in circAtlas v3.0, a comprehensive catalogue containing over 750,000 human circRNA ^15^ (**Table S3**). The LRDO enrichment and library preparation method detected a total of 69 circRNA by at least two reads, of which 5 were also detected by Ouro-seq (**Fig. 3A**). To explore the diversity of circRNA biogenesis we categorized circRNA based on their genomic localization compared to their gene of origin as annotated by the NCBI RefSeq assembly (see Methods). circRNA spanning a known exon were deemed “exonic”, and were the most abundant amongst plasma circRNA (154 exonic circRNA), while they formed a small proportion of the urine and CVS circRNA (20 and 31 exonic circRNA respectively). “Intronic” circRNA contain the introns of their gene of origin. In urine and CVS we detected 79 and 93 respectively, compared to only 9 intronic circRNA in plasma. “Antisense” circRNAs partially or fully overlap one or more annotated genes while originating from the opposite strand. Ouro-seq detected 53, 70 and 6 antisense circRNA in urine, CVS and plasma, respectively. Intriguingly, a large proportion of circRNA are from intergenic regions in urine and CVS (940 and 1,084 intergenic circRNA) compared to only 7 in plasma. This finding was confirmed by LRDO with 48 intergenic circRNA (∼90%) detected in both urine and CVS. Ten of these were found in both Ouro-seq and LRDO. Ouro-seq also detected a total of 29 circRNA mapping to unplaced scaffolds of the genome (**Fig. 3B**)(**Table S3**). The average size of exonic circRNA was 663.99 ± 513.57 bp, while intronic, intergenic and antisense circRNA were much longer with an average size of 1,319.49 ± 992.15 bp, 1,009.88 ± 732.77 bp and 1,175.06 ± 829.05 bp respectively (**Fig. 3C**).

**Figure 3.**
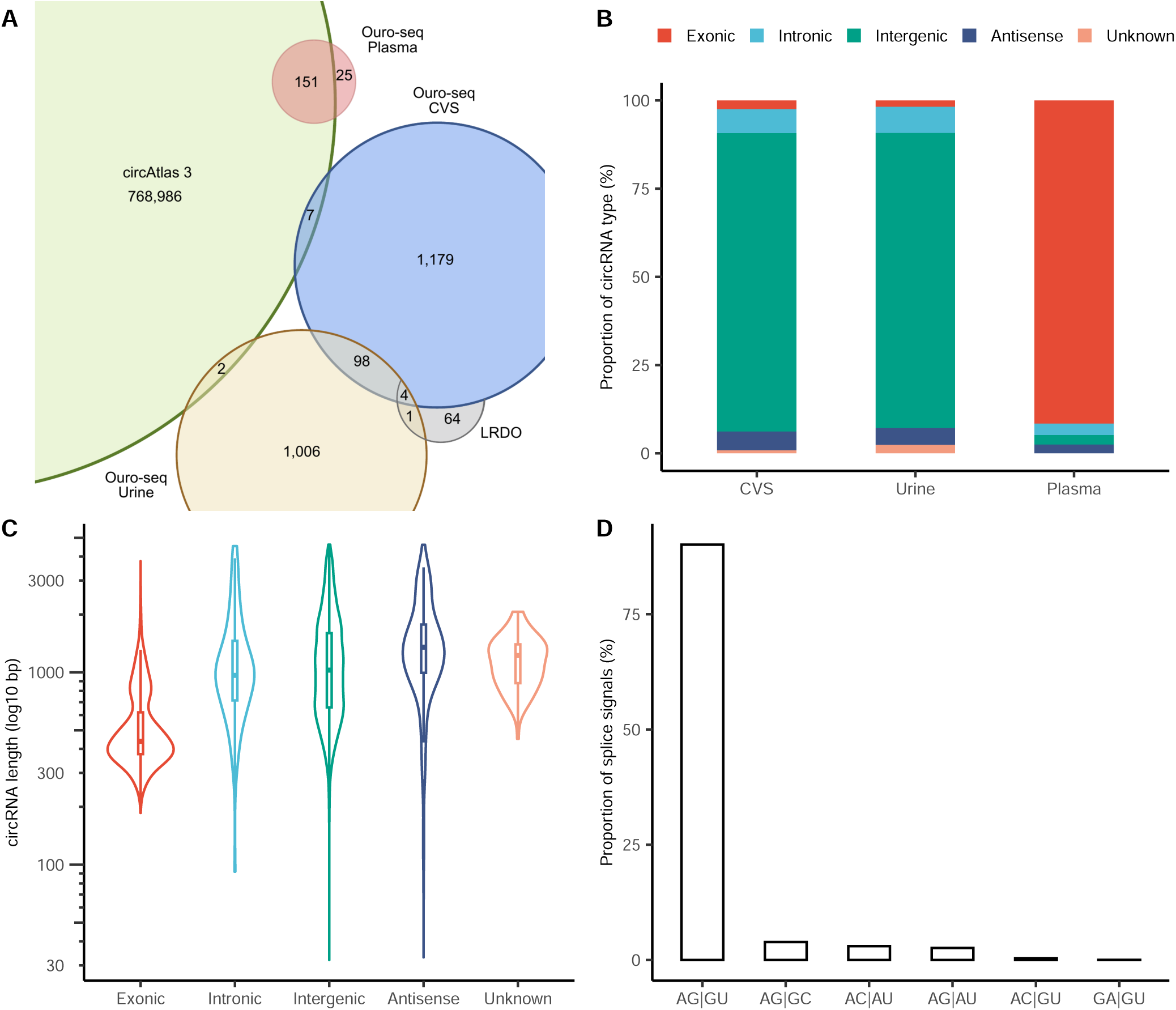
The hidden complexity of circRNA uncovered by Ouro-seq. **A.** The number of circRNA detected by Ouro-seq in CVS, urine and plasma compared to the number of circRNA detected by LRDO and present in the circAtlas 3 database. **B.** The proportion of circRNA types in different sample types detected by Ouro-seq. **C.** The length of full-length circRNA types detected by Ouro-seq. circRNA counts weighted with read counts. **D.** The proportion of splice signals flanking the BSJ.

A single gene locus can produce multiple circRNA either through variation of the back-splicing process, called alternative circularization, or through alternative splicing of the precursor mRNA. These processes are large contributors to circRNA diversity. Ouro-seq detected a total of 112 alternative circularization events, 46 for urine, 61 for CVS and 18 for plasma circRNA. Additionally, we detected 22 circRNA isoforms with alternative splicing: 13 in urine and 13 in CVS. 84% of the circRNA BSJs were flanked by the canonical AG/GU splice signal while the remaining circRNA bear alternative splice signals (**Fig. 3D**).

A recent study shows that transcriptional read-through events are remarkably common in the human transcriptome.^16^ circRNA from these read-through transcripts has been previously observed in tissue.^17^ Ouro-seq detected 42 read-through circRNA (rt-circRNA) with 26 overlapping two, 10 overlapping three and 6 overlapping more than 3 genes (**Table S3**). We detected 22 rt-circRNA from urine, 14 from CVS and 28 in plasma.

Although mitochondria lack the canonical splicing machinery, recent studies have described circRNA originating from the mitochondrial chromosome.^6,18^ We could also detect a total of 5 mitochondrial circRNA, 3 from CVS, one from urine and one from plasma (**Table S3**). Four of the 5 circRNA were transcribed from the light strand while one from the heavy strand of the mitochondrial genome. Two molecules are rt-circRNA, both overlapping two genes, one of them in antisense orientation.

These results show that Ouro-seq can detect a wide spectrum of various genomic and mitochondrial circRNA.

### Functional analysis of circRNA

circRNA can regulate gene expression through binding miRNAs also called miRNA sponging.^19^ We predicted a total of 6,759,211 miRNA binding sites, of which 510,361 were predicted by 3 tools (see Methods). A total of 283,698 of these sites were previously experimentally validated and 1,285,562 overlapped an AGO binding site. We accepted a miRNA binding site as a plausible site if it was experimentally validated, if it was predicted by at least 2 tools and overlapped an AGO binding site, or if it was detected by all 3 tools. This reduced the number to 997,409 plausible miRNA:circRNA interactions between 3,016 miRNA and 2,469 circRNA. Though many miRNA interactions do not show a canonical sequence complementarity (seed matching) between the miRNA and its target, seed matching, when present, can affect the targeting efficacy of the miRNA.^20^ Seed sequences for miRNA binding could not be detected for 47.68% of the interactions. Among detected seeds, the canonical 7mer-m8 represents 25.5% of interactions, while the rest of the seeds and seed mismatches contributed below 10%, with no distinct differences among circRNA types (**Fig. 4A**). The miRNA sponging capacity of circRNA might be limited by the stoichiometric imbalance between miRNA and circRNA. For effective sponging, circRNAs need either a relatively high abundance compared to miRNAs or many miRNA binding sites. The count of miRNA:circRNA interactions per kilobase showed a distinct pattern, with low interaction counts in intergenic circRNA and relatively high interaction counts in exonic and intronic circRNA (**Fig. 4B**). These results suggest a complex potential sponging specialization among circRNA types.

**Figure 4.**
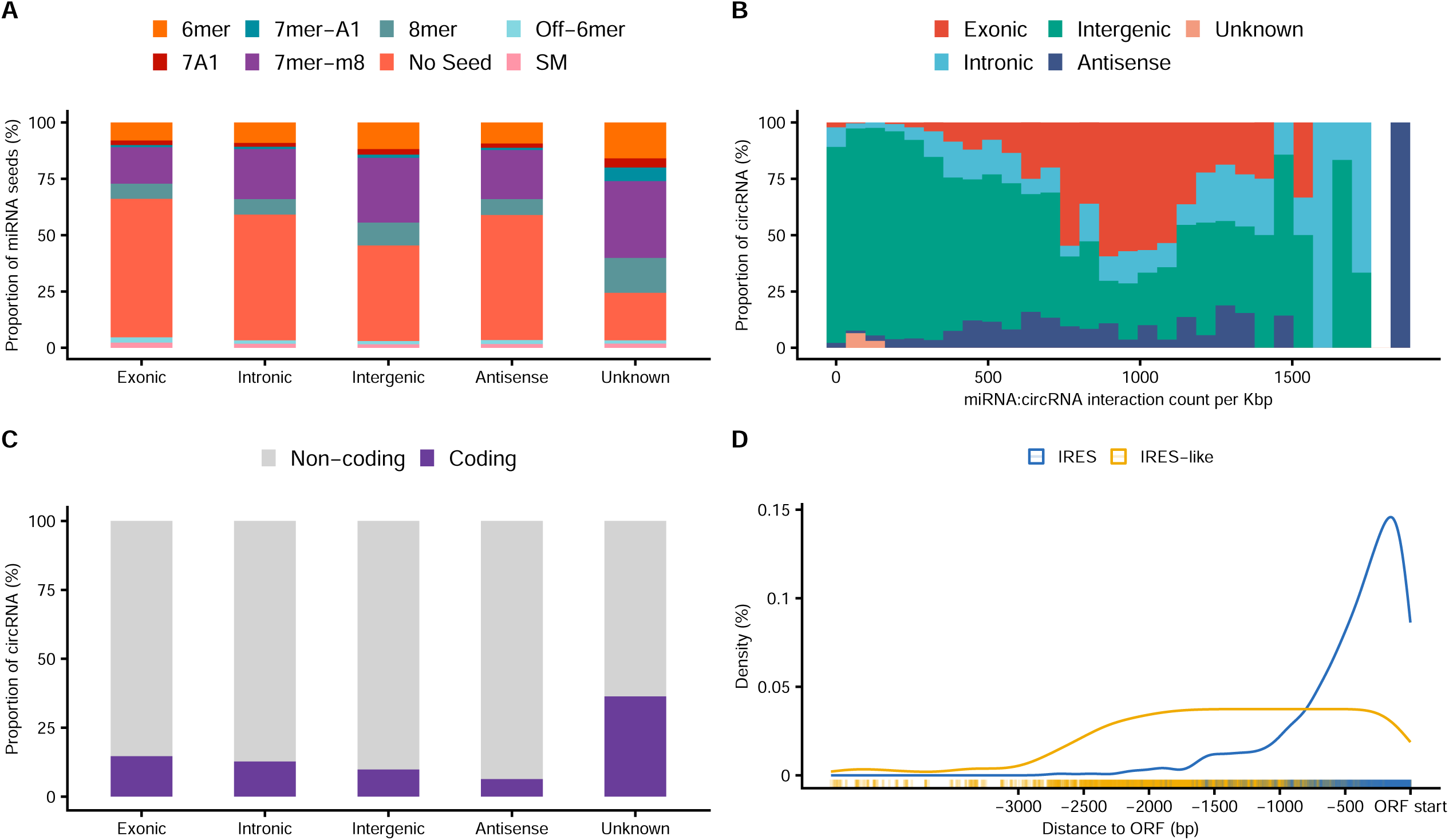
Functional analysis of circRNA. **A.** Proportion of miRNA seed types for circRNA categories. **B.** Cumulative proportion of circRNA types per miRNA:circRNA interaction counts. miRNA:circRNA interaction counts were normalized per Kbp and binned in 30 bins. **C.** The proportion of potentially coding and non-coding circRNA for each circRNA type. **D.** The density of IRES or IRES-like elements upstream a ORF start codon.

Recent studies uncovered that some circRNA might code for peptide or protein sequences, playing a role in tumorigenesis.^21–23^ We used CPC2 to predict the coding potential of circRNAs.^24^ CPC2 detected a total of 261 potentially coding circRNAs. CircRNA mapping to unplaced scaffolds had the highest coding potential (36.4% of circRNA), followed by exonic and intronic circRNA (14.7% and 12.8% of circRNA). In the absence of a 5’ cap, the translation machinery might access the open reading frames (ORFs) through binding to Internal Ribosomal Entry Sites (IRESs), or IRES-like hexamers.^25^ Using DeepIRES, we detected a total of 97 IRES upstream of a ORF start site predicted by CPC2 in 96 potentially coding circRNA. We also looked for IRES-like hexamers with translation initiation potential and found a total of 2,621 potential IRES-like hexamers upstream of an ORF start site predicted by CPC2 in 189 potentially circRNA. (**Fig. 4C**). The average distance of a predicted IRES from the start codon was 307.13 ± 388.74 bp, while IRES-like hexamers were located at an average 513.29 ± 563.12 bp upstream from a start codon (**Fig. 4D**).

## Discussion

Due to their scarcity in the total RNA pool, various methods have been developed to enrich circRNA for long-read sequencing. While effective with sufficient RNA input from tissues and cell cultures, their applicability to low-yield clinical biospecimens has been limited. Our study demonstrates the effectiveness of the Ouro-seq protocol for full-length circRNA enrichment and structural characterization from samples with limited RNA content such as liquid biopsies and cervico-vaginal self-samples. Full-length circRNA are present in low concentrations in liquid biopsies. The low total RNA input represents a substantial challenge for circRNA sequencing, particularly in plasma samples, where the input volume is also limited. In addition to linear RNA depletion and bead-based enrichment of long amplicons, Ouro-seq applies a short amplicon suppression PCR. By this further enrichment, our approach successfully identified a diverse array of circRNA molecules across different sample types, demonstrating the feasibility of liquid biopsy-based full-length circRNA analysis.

Our results challenge the established paradigm that exonic circRNA predominates in human samples. While previous literature has consistently reported exonic circRNA as the most abundant subtype in tissues and cell lines, we found this pattern only in plasma samples.^26^ In contrast, urine and CVS samples exhibited a strikingly different profile, with intergenic circRNA constituting the vast majority of detected molecules. This unexpected finding suggests that circRNA biogenesis and stability may be subject to yet undescribed tissue-specific or biofluid-specific regulatory mechanisms. The pronounced differences in circRNA types between plasma, urine, and CVS samples could reflect several biological phenomena. First, the cellular origin of circRNA in each biofluid likely differs, with plasma circRNA primarily originating from cells of the hematopoietic lineage, while urine and CVS circRNA may derive from bladder and cervico-vaginal epithelial cells.^27^ Second, the stability and half-life of different circRNA subtypes may vary across biofluids due to differences in preservatives, storage conditions, RNase activity, RNase sensitivity or exosome packaging.^2,28^

The canonical AG/GU splice signals predominated across all circRNA types, accounting for 84% of back-splice junctions, which aligns with the established understanding that circRNA biogenesis often leverages the canonical splicing machinery. Our analysis revealed that alternative splice signals are utilized more frequently in intergenic circRNA (16%) compared to intronic or exonic circRNA (12%). This observation suggests that intergenic circRNA may employ more diverse biogenesis mechanisms, potentially due to less constrained selection pressure on splicing signals in these regions, that warrant further investigation.

The relationship between circRNA length and miRNA interaction density revealed by our analyses provides compelling evidence for the role of circRNA in miRNA regulation. The relationship between circRNA size and miRNA binding site density suggests functional specialization, with certain circRNA potentially evolving to maximize their miRNA sponging capacity. This observation is further supported by our Specialization Potential Score analysis, which indicated that exonic circRNA, with their lower SPS scores, are particularly enriched for miRNA binding sites relative to peptide coding potential.

Furthermore, 10.6% of all detected circRNA were predicted as potentially coding, with 75% containing either a predicted IRES or one or more IRES-like hexamers upstream from the ORF. The translation of circRNAs into peptides is controversial, with some studies showing the biological prerequisites for translation initiation like IRES sequences or nucleotide modifications such as m^6^A being present on some circRNA.^29^ However, direct evidence for circRNA translation remains scarce.

Our detection of read-through circRNA (rt-circRNA) and mitochondrial circRNA further expands the known diversity of these molecules. The presence of rt-circRNA spanning multiple genes suggests complex transcriptional events that may contribute to regulatory networks involving multiple genomic loci. Similarly, the detection of mitochondrial circRNA, despite the absence of canonical splicing machinery in mitochondria, points to alternative mechanisms of circRNA biogenesis that merit further exploration.

## Limitations of Study

Our results are limited by the relatively small sample size. This small cohort precludes robust statistical analyses and limits our ability to draw definitive conclusions about the diagnostic or prognostic potential of circRNA profiles. Future studies should incorporate larger cohorts, and more robust controls, such as xenograft models of the specific cancer types, or matched tissue and liquid biopsy samples for both conditions. Such controls would validate the identified circRNA signatures and provide insights into their biological relevance and clinical applicability.

Second, urine samples have a relatively high total RNA concentration compared to plasma, which suggests contribution of cellular RNA. While this can improve the detection of certain cancer types, like bladder cancer that sheds directly into the lumen of the bladder or in our case cervical cancer, where the tumor-derived RNA can get into the urine through vaginal secretions, for most cancer types the tumor-derived cell-free circRNA signal will be diluted by the cellular signal. For these cancer types, the use of urine supernatant after centrifugation may improve detection.^30^

Third, artefactual BSJs might result from template switching (TS) events during reverse transcription. This was shown to affect circRNA detection using SRS, for which the detection is based only on the presence of BSJs.^31^ We based the detection of circRNA on two criteria: the concatameric nature of the reads and the presence of canonical or alternative splice signals flanking the BSJ. Further evidence is needed to conclude if these stringent criteria together with the elevated reverse transcription temperatures of Ouro-seq reduce the possibility of false circRNA detection.

Fourth, accurate quantification of such low-abundance molecules by sequencing is inherently challenging due to stochastic variation in molecular sampling. While more sensitive quantification methods such as qRT-PCR, ddPCR, or microarrays exist, these approaches require *a priori* isoform-level sequence information that can only be obtained through sequencing-based approaches. Consequently, we emphasize Ouro-seq’s utility for discovering and characterizing novel circRNA isoforms rather than for precise quantification.

Fifth, current circRNA detection tools, including those employed in our study, may have suboptimal sensitivity and specificity.^32^ The computational identification of circRNA remains challenging due to the sequence similarity between circRNA and their linear counterparts, as well as the complexity of back-splice junction detection. Improvements in bioinformatics approaches, potentially leveraging machine learning algorithms, could enhance the accuracy of circRNA identification and characterization.

Sixt, while our study provides insights into the potential functional roles of different circRNA types, experimental validation is needed to confirm these predictions. Future work should include functional assays to directly assess the miRNA sponging capacity of candidate circRNA and to verify the translation of predicted coding sequences.

Finally, the biological significance of the diverse circRNA population we detected, particularly the abundant intergenic circRNA in urine and CVS, remains to be elucidated. Comprehensive analyses of circRNA expression across diverse tissue types and disease states will be essential to understand the physiological and pathological roles of these molecules.

## Conclusions

Our study introduces Ouro-seq, a novel protocol that improves the recovery and characterization of full-length circRNA from samples with reduced RNA content. The unexpected predominance of intergenic circRNA in urine and CVS samples challenges existing paradigms and highlights the need for comprehensive analyses across different biofluid types. The associations between circRNA characteristics and their potential functions provide insights into specialized roles in miRNA regulation and possible peptide coding capacity. Despite limitations in sample size, this work expands the current understanding of circRNA biology in liquid biopsies and provides a foundation for future investigations into their diagnostic and therapeutic potential.

## Acknowledgements

The authors thank the Amsterdam UMC Liquid Biopsy Center and Amsterdam UMC Core Facility Genomics for logistical support. The authors are also thankful to Mengfei Xu and Mariano A. Molina Beitia for their comments on RNA extraction optimization and Justus Olijslager and Frederieke Koller for their suggestions on the ONT protocol and sequencing optimization. B.W. was supported by the Stichting NEXTGEN HIGHTECH Program (Biomed02). N.M. and Y.v.d.B. were supported by The Netherlands Organisation for Health Research and Development (ZonMW) Off-road grant [04510012210008]. F.M. and N.M. were supported by a Dutch Cancer Fund (KWF14976 and KWF14450). The SOLUTION1 study was financially supported by the Hanarth Foundation and Weijerhorst Foundation. This work was carried out on the Dutch national e-infrastructure with the support of SURF Cooperative. Funders have no role in the design of the study.

## Author contributions

Conceptualization and design, N.M., B.W., F.M. and R.D.M.S.; experiments and data collection, B.W. and Y.v.d.B.; data processing, N.M.; software development, N.M.; data analysis, N.M.; sample acquisition, M.P. and M.B.; funding acquisition, N.M., R.D.M.S., M.B. and F.M.; manuscript draft, N.M., B.W. and Y.v.d.B.; manuscript revision and comments, N.M., B.W., Y.v.d.B., F.M., M.P., M.B. and R.D.M.S.; supervision, N.M.

## Supplementary information

**Supplementary information 1: The oligonucleotides used in this study.** P-SSP: Panhandle Strand Switching Primer, V is a randomly incorporated A, C or G and the [GGG]s are 2’

O-methyl RNA Gs; RH-P-CRTA: Random-Hexamer Panhandle Adapter, N is a randomly incorporated A, C, G or T; P-PCR: Panhandle PCR primer.

**Table S1: Sample types and concentrations.** Total RNA represents the RNA content of the samples after RNA extraction while Depleted RNA is the RNA concentration after ribosomal and linear RNA depletion. RNA concentration was measured using the Qubit™ RNA HS Assay Kit and fluorometer.

**Table S2: Read statistics of the samples.** CVS: cervico-vaginal self-sample, LRDO: linear RNA degradation only protocol.

**Table S3: The detected circRNAs.** Isoforms are shown as intervals of exons separated by a coma. Splice signals represent the signals flanking the back-splice junction represented by |. circRNA with a list of gene names are read-through circRNA. CVS: cervico-vaginal self-sample, LRDO: linear RNA degradation only protocol.

## Methods

### Study population

Ouro-seq was applied to a total of 22 clinical samples collected from 19 individuals. Samples included urine (n = 5) and CVS (n = 5) from cervical cancer patients (n = 7) and plasma (n = 12) from lung cancer patients (n = 7) and healthy volunteers without known malignancy (n = 5) (**Table S1**). Samples from cancer patients were collected before treatment. Cervical cancer patients were enrolled in the SOLUTION studies at tertiary care medical centers.^33^ Lung cancer patients and healthy volunteers were recruited through the Liquid Biopsy Center at Amsterdam UMC, location VUmc.^34^

This study was conducted in compliance with the Declaration of Helsinki. All participants were 18 years or older and provided written informed consent prior to sample collection. Ethical approval was provided by the Medical Ethical Committee of Amsterdam UMC for the use of samples collected through the SOLUTION studies (METc 2016.213 and METc 2022.0538; Trial registration ID NL56664.029.16) and Amsterdam UMC Liquid Biopsy Center (METc U2019_035). SOLUTION study samples were biobanked under METc 2022.0819.

### Clinical sample collection and processing

Blood samples were collected in EDTA-K2 tubes and processed via double centrifugation (900g for 7 min, followed by 2,500g for 10 min at room temperature). Plasma supernatant was divided into 0.5 ml Nunc tubes and stored at -80°C. Urine and CVS were collected at home, mailed and further processed at the pathology department of Amsterdam UMC, location VUmc, as previously described.^33^ In short, urine was collected using 30 mL collection tubes containing 40 mM EDTA. Full void urine (*i.e.* unfractionated urine) was stored at -20°C (SOLUTION1) or -80°C (SOLUTION Time). CVS were collected using a dry-brush device (Evalyn Brush, Rovers Medical Devices, Oss, The Netherlands), which was placed in 1.5 mL ThinPrep PreservCyt medium (Hologic, Marlborough, MA, USA) upon arrival and stored at 4°C.

### Total RNA extraction

Total RNA was extracted from plasma (5 mL), full void urine (10 mL), and CVS (200 µL) using the Plasma/Serum RNA Purification Kit (Norgen Biotek Corp, Ontario, Canada), Urine Total RNA Purification Maxi Kit Dx (Slurry Format; Norgen) and TRIzol Reagent (Thermo Fisher Scientific, Waltham, MA, USA), respectively, following manufacturers’ guidelines. For urine RNA extractions, 100 µL of 3M Sodium Acetate (pH 5.2) was added as RNA carrier. Total RNA was quantified using the Qubit™ RNA BR Assay Kit (Invitrogen, Carlsbad, CA, USA) or the NanoDrop 1000 (Thermo Fisher) for low-yield samples.

### circRNA enrichment

circRNA enrichment was performed by ribosomal RNA (rRNA) depletion and subsequent linear RNA degradation. For each sample, up to 2,660 ng total RNA was subjected to rRNA depletion using the RiboMinus™ Eukaryote System v2 (Thermo Fisher Scientific), following manufacturers’ guidelines. A total of 5 ng synthetic circRNA spike-in (Creative Biogene) was added to each sample prior to rRNA depletion for normalization and internal control purposes (**Supplementary information 1**). Briefly, rRNA molecules were selectively hybridized with the RiboMinus™ Eukaryote Probe Mix v2 (Thermo Fisher Scientific). Resulting rRNA-probe complexes were captured and removed through RiboMinus™ magnetic bead-based separation. rRNA-depleted RNA was purified using Nucleic Acid Binding Beads (Thermo Fisher Scientific) and quantified using the Qubit™ RNA BR Assay Kit (Invitrogen).

Next, linear RNA fragments were selectively removed by poly(A) tailing and subsequent RNAse R treatment. Poly(A) tailing was carried out by incubating rRNA-depleted RNA in a 20 µL reaction mixture containing 2 µL *E. coli* Poly(A) Polymerase (5 U/µL; New England Biolabs, Ipswich, MA, USA), 2 µL 10x *E. coli* Poly(A) Polymerase Reaction Buffer (NEB), and 2 µL ATP (10 mM; NEB) at 37°C for 30 min. Polyadenylated RNA was purified using Agencourt RNAClean XP beads (Beckman Coulter, Brea, CA, USA) following manufacturers’ guidelines. Next, polyadenylated RNA was treated with 10 U RNAse R (20 U/µL; Epicentre, Madison, WI, USA) in a 20 µL reaction containing 1x RNAse R Reaction buffer and incubated at 37°C for 15 min. RNAse R treated RNA was cleaned using Agencourt RNAClean XP beads (Beckman Coulter) and quantified using the HS RNA ScreenTape assay (Agilent Technologies, Santa Clara, CA, USA) on an Agilent 4200 TapeStation for quality control before library preparation.

### Library preparation and sequencing

Total enriched circRNA from urine and CVS samples was evenly divided into two aliquots and underwent library preparation following both LRDO and our customized Ouro-seq approach for Oxford Nanopore Technologies (ONT) long-read cDNA-PCR sequencing, allowing for a head-to-head comparison of both workflows. Enriched circRNA from plasma samples only underwent Ouro-seq due to limited circRNA yields recovered from this sample type.

For LRDO, samples were processed using the ONT SQK-PCB114.24 workflow following manufacturers’ instructions, including steps adopted from the sequence-specific SQK-PCB111 protocol. In short, rolling circle reverse transcription was carried out by incubating up to 219 ng enriched circRNA with 1 µL RH-CRT adapter (2 µM; Eurogentec) and 1 µL dNTPs (10 mM; NEB) in a total reaction volume of 11 µL at 65°C for 5 min. After snap-cooling for 1 min, 4 µL Maxima H Minus 5x RT Buffer (Thermo Fisher) and 2 µL Strand Switching Primer II (SSPII; ONT) were added to the reaction mixture with a total volume of 19 µL and incubated at 35°C for 2 min. Then, 200 U of Maxima H Minus Reverse Transcriptase (200 U/μL; Thermo Fisher) was added and reverse transcription was carried out at 42°C for 90 min followed by 85°C for 5 min. cDNA PCR amplification and barcoding was performed using the ONT SQK-PCB114.24 kit according to standard manufacturers’ guidelines with 30 PCR cycles. To dilute reverse transcriptase prior to PCR, cDNA was divided into four 5 µL reactions. Libraries were purified and size selected using 0.5X AMPure XP Beads (Beckman Coulter).

For Ouro-seq, libraries were enriched for longer transcripts by adapting a previously described workflow.^35^ Firstly, rolling circle reverse transcription was carried out using a custom cDNA RT adapter and Panhandle Strand Switching Primer (**Supplementary information 1**). Up to 219 ng circRNA was incubated with 1 µL Random Hexamer Panhandle adapter (RH-P-CRT 10 µM; Eurogentec) (**Supplementary information 1**) and 1 µL dNTPs (10 mM; NEB) in a total reaction volume of 11 µL at 65°C for 5 min. After snap-cooling for 1 min, 4 µL Maxima H Minus 5x RT Buffer, 2 µL Betaine (5M; Sigma-Aldrich, St. Louis, MO, USA) and 2 µL Panhandle Strand Switching Primer (P-SSP, 10 µM; Eurogentec) were added to the reaction mixture with a total volume of 19 µL and incubated at 35°C for 2 min. Next, 200 U of Maxima H Minus Reverse Transcriptase (200 U/μL; Thermo Fisher) was added and reverse transcription was performed using the following cycling conditions: 50°C for 30 min, by 55°C for 5 min, 50°C for 5 min, 60°C for 1 min, 55°C for 3 min, 50°C for 3 min, 65°C for 1 min, 60°C for 3 min, 50°C for 3 min, 75°C for 1 min, 70°C for 3 min, 50°C for 3 min, 85°C for 10 min. Secondly, cDNA PCR amplification with short-molecule PCR suppression was performed to enrich for longer transcripts. cDNA was split into four reactions of 5 µL each to dilute reverse transcriptase prior to PCR. Per PCR reaction, 0.5 µL of Panhandle primer (P-PCR, 10 µM; Eurogentec) (**Supplementary information 1**) and 1x LongAmp Hot Start Taq Master Mix (NEB) were added in a total reaction volume of 25 µL. PCR was carried out using the following cycling conditions: initial denaturation at 95°C for 1 min, followed by 5 cycles of 95°C for 20 s, 58°C for 4 min, and 68°C for 6 min, then 30 cycles of 95°C for 20 s, 64°C for 30 s, and 68°C for 5 min, with a final extension at 72°C for 5 min. After PCR, excess primers were removed by incubation with Exonuclease I (NEB) for 15 min at 37°C followed by 15 min at 80°C. Libraries were purified and size selected using 0.5X AMPure XP Beads (Beckman Coulter). Lastly, Ouro-seq libraries underwent End-repair and dA tailing using the Ultra II End Prep / dA tailing Module (NEB) and native barcode ligation using the ONT Native Barcoding Kit 24 V14 (SQK-NBD114.24) following standard manufacturers’ instructions.

Both conventional LRDO cDNA-seq and Ouro-seq libraries were quantified using the D5000 ScreenTape assay on an Agilent 4200 TapeStation for quality control before pooling. Barcoded libraries were pooled in equimolar amounts and adapter ligated using rapid adapters for LRDO (SQK-PCB114.24) and native adapters for Ouro-seq (SQK-NBD114.24). Long-read sequencing was performed using a R10.4.1 flow cell on a PromethION platform (ONT).

### Data preprocessing

Reads were base called with Dorado v.0.7.2 [https://github.com/nanoporetech/dorado] using model dna_r10.4.1_e8.2_400bps_sup@v4.3.0 with modification detection turned off. Data pre-processing and analysis was performed using our Ouro-tools Snakemake v.8.18.0 [https://github.com/snakemake] pipeline [https://github.com/moldovannorbert/Ouro-tools] as follows: Base-called reads were trimmed using Porechop v.0.2.4 [https://github.com/rrwick/Porechop] with *--extra_end_trim 0* parameter to avoid trimming of splice signals around the BSJ. For quality check and read length extraction trimmed reads were aligned to GRCh38 genomic reference [GeneBank assembly: GCA_000001405.28] complemented with the sequence of a synthetic circularized RNA spike-in (**Supplementary information 1**) as a separate scaffold using minimap2 v.2.26 [https://github.com/lh3/minimap2] with the *-ax splice* parameter. Samtools v.1.17 [https://github.com/samtools/samtools] was used to sort and filter reads with parameters *-q 5 -F 4 -F 256 -F 1024 -F 2048*. NanoPlot v.1.41.6 [https://github.com/wdecoster/NanoPlot] was used for QC statistics with the *--alength --huge* parameters.

### circRNA calling and isoform collapse

circRNA candidate identification was performed on each sample using the Ouro-tools implementation of CIRI-long v.1.1.0 [https://github.com/bioinfo-biols/CIRI-long]. In brief, for circRNA identification we ran *CIRI-long call* with the GRCh38 genomic reference complemented with the sequence of the synthetic circularized RNA spike-in as a separate scaffold and NCBI RefSeq genome assembly v.2022-10-28. Isoforms were identified, counted and circRNA types were determined using *CIRI-long collapse* using the same genomic reference and assembly as for circRNA calling.

### Database comparison

We compared the BSJ coordinates of circRNAs identified in this study with those documented in the circAtlas v3.0 human_bed_v3 database (hg38 version, retrieved October 18, 2024). CircRNAs were classified as previously described if their BSJ coordinates matched entries in the database within a ±5 bp window.

### miRNA:circRNA association prediction

We used Circr commit 22a50bb [https://github.com/bicciatolab/Circr] for detecting miRNA binding sites on the full length of predicted circRNA. In brief, Circr combines miRanda v.3.3a, RNAhybrid v2.1.2 and TargetScan v.7.0 for *de novo* miRNA binding site prediction, followed by annotation with experimentally validated miRNA:RNA interactions and Argonaut (AGO) target sites. We ran Circr with the parameters *-c -s human -v hg38*, the GRCh38 genomic reference, the NCBI RefSeq genome assembly v.2022-10-28 for gene annotation and files provided with the tool for rRNA coordinates, miRNA sequences, experimentally validated miRNA:RNA interactions and AGO annotations.

### Coding potential computation

We used the standalone version of *CPC2 v.1.0.1* [https://github.com/gao-lab/CPC2_standalone] with default settings to classify the circRNA’s coding potential. CPC2 uses the Fickett score, the length, the integrity and the isoelectric point calculated based on the longest putative ORF per circRNA to classify molecules using a support vector machine classifier trained on high-confidence human coding and non-coding transcripts.^24^ CircRNA with a coding probability of 0.5 or higher were deemed “coding” while those below 0.5 “non-coding”. To detect ORFs overlapping the BSJ we performed the ORF calling in an extended circRNA sequence by concatenating two copies of the same sequence.

### IRES prediction

We ported DeepIRES [https://github.com/SongLab-at-NUAA/DeepIRES] to Linux by changing operation system specific dependencies and paths. DeepIRES is a hybrid deep learning tool incorporating dilated 1D convolutional neural network blocks, bidirectional gated recurrent units, and a self-attention module for improved IRES prediction.^36^ Additionally, we searched the circRNA sequences for 97 IRES-like hexamers retrieved from a previous publication.^25^ We called a motif predicted IRES or predicted IRES-like if located upstream the start of an ORF predicted by CPC2.

### Statistics and figures

All statistical analyses were performed in *R* v.4.5.0 [https://www.r-project.org/]. Data processing and summary statistics were conducted using the *dplyr v.1.1.4*, *tidyr v.1.3.1*, and *stringr v.1.5.1* packages. For summary statistics we computed the mean and the standard deviation as mean ± SD. Statistical significance was evaluated using the Mann-Whitney U test with an alpha value of 0.05. Figures were generated in *R* using *ggplot2 v.3.5.2 ggpubr v.0.6.0*, *ggsci v.3.2.0* and *magick v.2.8.6*. Schematic representation of the Ouro-seq protocol and the experiments were created using *Biorender*. Figures were arranged and exported as multi-panel PDFs with consistent axis scaling, color schemes, and annotation styles.

## References

1. Verduci, L., Tarcitano, E., Strano, S., Yarden, Y., and Blandino, G. (2021). CircRNAs: role in human diseases and potential use as biomarkers. Cell Death Dis. 12, 468.

2. Jeck, W.R., and Sharpless, N.E. (2014). Detecting and characterizing circular RNAs. Nat. Biotechnol. 32, 453–461.

3. Enuka, Y., Lauriola, M., Feldman, M.E., Sas-Chen, A., Ulitsky, I., and Yarden, Y. (2016). Circular RNAs are long-lived and display only minimal early alterations in response to a growth factor. Nucleic Acids Res. 44, 1370–1383.

4. Wang, C., and Liu, H. (2022). Factors influencing degradation kinetics of mRNAs and half-lives of microRNAs, circRNAs, lncRNAs in blood in vitro using quantitative PCR. Sci. Rep. 12, 7259.

5. Steijger, T., Abril, J.F., Engström, P.G., Kokocinski, F., RGASP Consortium, Hubbard, T.J., Guigó, R., Harrow, J., and Bertone, P. (2013). Assessment of transcript reconstruction methods for RNA-seq. Nat. Methods 10, 1177–1184.

6. Zhang, J., Hou, L., Zuo, Z., Ji, P., Zhang, X., Xue, Y., and Zhao, F. (2021). Comprehensive profiling of circular RNAs with nanopore sequencing and CIRI-long. Nat. Biotechnol. 39, 836–845.

7. Gao, Y., Zhang, J., and Zhao, F. (2018). Circular RNA identification based on multiple seed matching. Brief. Bioinform. 19, 803–810.

8. de Jong, L.C., Cree, S., Lattimore, V., Wiggins, G.A.R., Spurdle, A.B., kConFab Investigators, Miller, A., Kennedy, M.A., and Walker, L.C. (2017). Nanopore sequencing of full-length BRCA1 mRNA transcripts reveals co-occurrence of known exon skipping events. Breast Cancer Res. 19, 127.

9. Kiyose, H., Nakagawa, H., Ono, A., Aikata, H., Ueno, M., Hayami, S., Yamaue, H., Chayama, K., Shimada, M., Wong, J.H., et al. (2022). Comprehensive analysis of full-length transcripts reveals novel splicing abnormalities and oncogenic transcripts in liver cancer. PLoS Genet. 18, e1010342.

10. Hou, L., Zhang, J., and Zhao, F. (2023). Full-length circular RNA profiling by nanopore sequencing with CIRI-long. Nat. Protoc. 18, 1795–1813.

11. Xin, R., Gao, Y., Gao, Y., Wang, R., Kadash-Edmondson, K.E., Liu, B., Wang, Y., Lin, L., and Xing, Y. (2021). isoCirc catalogs full-length circular RNA isoforms in human transcriptomes. Nat. Commun. 12, 266.

12. Rahimi, K., Venø, M.T., Dupont, D.M., and Kjems, J. (2021). Nanopore sequencing of brain-derived full-length circRNAs reveals circRNA-specific exon usage, intron retention and microexons. Nat. Commun. 12, 4825.

13. Fuchs, S., Babin, L., Andraos, E., Bessiere, C., Willier, S., Schulte, J.H., Gaspin, C., and Meggetto, F. (2022). Generation of full-length circular RNA libraries for Oxford Nanopore long-read sequencing. PLoS One 17, e0273253.

14. Cabús, L., Lagarde, J., Curado, J., Lizano, E., and Pérez-Boza, J. (2022). Current challenges and best practices for cell-free long RNA biomarker discovery. Biomark Res 10, 62.

15. Wu, W., Ji, P., and Zhao, F. (2020). CircAtlas: an integrated resource of one million highly accurate circular RNAs from 1070 vertebrate transcriptomes. Genome Biol 21, 101.

16. Caldas, P., Luz, M., Baseggio, S., Andrade, R., Sobral, D., and Grosso, A.R. (2024). Transcription readthrough is prevalent in healthy human tissues and associated with inherent genomic features. Commun Biol 7, 100.

17. Vo, J.N., Cieslik, M., Zhang, Y., Shukla, S., Xiao, L., Zhang, Y., Wu, Y.-M., Dhanasekaran, S.M., Engelke, C.G., Cao, X., et al. (2019). The Landscape of Circular RNA in Cancer. Cell 176, 869–881.e13.

18. Wu, Z., Sun, H., Wang, C., Liu, W., Liu, M., Zhu, Y., Xu, W., Jin, H., and Li, J. (2020). Mitochondrial Genome-Derived circRNA mc-COX2 Functions as an Oncogene in Chronic Lymphocytic Leukemia. Mol Ther Nucleic Acids 20, 801–811.

19. Hansen, T.B., Jensen, T.I., Clausen, B.H., Bramsen, J.B., Finsen, B., Damgaard, C.K., and Kjems, J. (2013). Natural RNA circles function as efficient microRNA sponges. Nature 495, 384–388.

20. Bartel, D.P. (2009). MicroRNAs: target recognition and regulatory functions. Cell 136, 215–233.

21. Meng, E., Deng, J., Jiang, R., and Wu, H. (2022). CircRNA-Encoded Peptides or Proteins as New Players in Digestive System Neoplasms. Front Oncol 12, 944159.

22. Zheng, X., Chen, L., Zhou, Y., Wang, Q., Zheng, Z., Xu, B., Wu, C., Zhou, Q., Hu, W., Wu, C., et al. (2019). A novel protein encoded by a circular RNA circPPP1R12A promotes tumor pathogenesis and metastasis of colon cancer via Hippo-YAP signaling. Mol Cancer 18, 47.

23. Yang, Y., Gao, X., Zhang, M., Yan, S., Sun, C., Xiao, F., Huang, N., Yang, X., Zhao, K., Zhou, H., et al. (2018). Novel Role of FBXW7 Circular RNA in Repressing Glioma Tumorigenesis. J Natl Cancer Inst 110, 304–315.

24. Kang, Y.-J., Yang, D.-C., Kong, L., Hou, M., Meng, Y.-Q., Wei, L., and Gao, G. (2017). CPC2: a fast and accurate coding potential calculator based on sequence intrinsic features. Nucleic Acids Res 45, W12–W16.

25. Fan, X., Yang, Y., Chen, C., and Wang, Z. (2022). Pervasive translation of circular RNAs driven by short IRES-like elements. Nat Commun 13, 3751.

26. Pisignano, G., Michael, D.C., Visal, T.H., Pirlog, R., Ladomery, M., and Calin, G.A. (2023). Going circular: history, present, and future of circRNAs in cancer. Oncogene 42, 2783–2800.

27. Memczak, S., Papavasileiou, P., Peters, O., and Rajewsky, N. (2015). Identification and Characterization of Circular RNAs As a New Class of Putative Biomarkers in Human Blood. PLoS One 10, e0141214.

28. Xu, Y., Kong, S., Qin, S., Shen, X., and Ju, S. (2020). Exosomal circRNAs: Sorting Mechanisms, Roles and Clinical Applications in Tumors. Front Cell Dev Biol 8, 581558.

29. Yang, Y., Fan, X., Mao, M., Song, X., Wu, P., Zhang, Y., Jin, Y., Yang, Y., Chen, L.-L., Wang, Y., et al. (2017). Extensive translation of circular RNAs driven by N-methyladenosine. Cell Res 27, 626–641.

30. Peng, H., Pan, M., Zhou, Z., Chen, C., Xing, X., Cheng, S., Zhang, S., Zheng, H., and Qian, K. (2024). The impact of preanalytical variables on the analysis of cell-free DNA from blood and urine samples. Front Cell Dev Biol 12, 1385041.

31. Tang, C., Yu, T., Xie, Y., Wang, Z., McSwiggin, H., Zhang, Y., Zheng, H., and Yan, W. (2018). Template switching causes artificial junction formation and false identification of circular RNAs. bioRxiv. 10.1101/259556.

32. Vromman, M., Anckaert, J., Bortoluzzi, S., Buratin, A., Chen, C.-Y., Chu, Q., Chuang, T.-J., Dehghannasiri, R., Dieterich, C., Dong, X., et al. (2023). Large-scale benchmarking of circRNA detection tools reveals large differences in sensitivity but not in precision. Nat Methods 20, 1159–1169.

33. van den Helder, R., Steenbergen, R.D.M., van Splunter, A.P., Mom, C.H., Tjiong, M.Y., Martin, I., Rosier-van Dunné, F.M.F., van der Avoort, I.A.M., Bleeker, M.C.G., and van Trommel, N.E. (2022). HPV and DNA Methylation Testing in Urine for Cervical Intraepithelial Neoplasia and Cervical Cancer Detection. Clin Cancer Res 28, 2061–2068.

34. van der Pol, Y., Tantyo, N.A., Evander, N., Hentschel, A.E., Wever, B.M., Ramaker, J., Bootsma, S., Fransen, M.F., Lenos, K.J., Vermeulen, L., et al. (2023). Real-time analysis of the cancer genome and fragmentome from plasma and urine cell-free DNA using nanopore sequencing. EMBO Mol Med 15, e17282.

35. Bayega, A., Oikonomopoulos, S., Wang, Y.C., and Ragoussis, J. (2022). Improved Nanopore full-length cDNA sequencing by PCR-suppression. Front Genet 13, 1031355.

36. Zhao, J., Chen, Z., Zhang, M., Zou, L., He, S., Liu, J., Wang, Q., Song, X., and Wu, J. (2024). DeepIRES: a hybrid deep learning model for accurate identification of internal ribosome entry sites in cellular and viral mRNAs. Brief Bioinform 25. 10.1093/bib/bbae439.

