## Supplementary information 1 for "Ouro-seq: Improved Recovery of Full-Length circRNAs from Samples with Limited RNA Content"

>circRNA_spike_in
ACGGAGGAGCTTTGGCATACTAGGCTAGCGAATCTGCAACTAACGCAAGTTACATCCTAGCTAG
CGAAGGGCGTCCCAATTTTCGCTAACCCGACGCGACGCATAAAAAGCGAGAATAACGCCTAAG
GGATGTACAATGGATGTTGATTATGCCTTCGGGAATGAGGGATGATTTGCGAAAAACAAGTCAA
TACCTAACCAAATCCGCTAATGGACACCCGTAATCGTGCCCAAGTTTAACTGGTCGGTAGGTGG
CAGGCAAAGCGCTAGTATCCCTAGGCGCGACACTATAAGTTTACAACTGCGAGAATTGACACTA
TGAGCGCGCATACTGGGGCCAGAATAGGCAATACCATGTGCGTCCCTGTGTGAACAGCTCGC
GGCCATCAGAAGTTGGGATTGACGCATGATCTTGATCGAGCATACGGCTTCCACCAACCCATA
GTACTTGGTAACTATAGCAATCAAGCACGCGTGAGCACAACGCTATCCAAATTACTACATTAACT
GG

>P-SSP
AAGCAGTGGTATCAACGCAGAGTGGTTTVVVVTTVVVVTTVVVVTTVVVVTTT[GGG]

>RH-P-CRTA
AAGCAGTGGTATCAACGCAGAGTATGCAACGCAACTNNNNNN

>P-PCR
TCGTCGGCAGCGTCAAGCAGTGGTATCAACGCAGAGT
